## Supplementary material for "Hierarchical dinucleotide distribution in genome along evolution and its effect on chromatin packing": SI

**2.Biomedical Pioneering Innovation Center (BIOPIC), Peking University, Beijing 100871, China**

**3.Beijing Advanced Innovation Center for Genomics (ICG), Peking University, Beijing 100871, China**

**I Supplementary Text**

**II Supplementary Tables**

**III Supplementary Figures**

### Supplementary Text

#### Data sources

Genomes of species are listed in Table S1. They were retrieved from databases at UCSC genome browser and NCBI. For each species, the longest chromosome was chosen to calculate the Variability, Multi-scale entropy (SE) and Pearson correlation coefficient of the dinucleotide density. Source and information of Hi-C datasets are listed in Table S5. Source of RNA-seq are listed in Table S6. Cell types of Hi-C map are listed in Table S7.

#### Dinucleotide density calculation

The dinucleotide density was calculated along the genome with a non-overlapped sliding window. The window size was chosen according to the genome length scale interested. To calculate the variability of the dinucleotide density, the sliding window size was chosen as 1000bp, comparable to the length of the CpG island. To calculate the Pearson correlation coefficient of the dinucleotide density, the sliding window size was chosen as 200bp.

**Decomposing dinucleotides density series by Hilbert Huang Transform (HHT)**

To analyze the dinucleotide distribution at local scale, the dinucleotide density series need to be decomposed into fluctuation series at different frequencies. Hilbert Huang Transform (HHT) can decompose a data series into oscillatory modes of different frequencies[1], which is an adaptive method can be applied to the nonlinear and nonstationary process. Unlike Fourier Transformation, HHT is not limited by the Heisenberg uncertainty principle and does not require a priori basis, which enables HHT very suitable to decompose and analyze the density series with lots of drastic fluctuations.

Take the CpG density series of human and Nile Tilapia as example, they can be decomposed to series at different frequencies by HHT (Figure S1), respectively. Their decomposed series at high frequency are distinct from each other. Sharp peaks exist in the curve of decomposed series of human, while they are absent in the ones of Nile Tilapia, which leads to a larger CpG Variability of human than the one of Nile Tilapia.

**Validation of 3σ threshold in calculating the variability of dinucleotide density**

To calculate the variability, the absolute values of decomposed density series were divided into the high amplitude group in which the values are larger than a+3σ, and the low amplitude group with values smaller than a+σ, where σ is the standard deviation and a is the average of the absolute values. Threshold value 3σ is chosen for it is often used as a threshold in detecting outliers in the statistics theory. The CpG variability was also calculated using the threshold value 2σ, and the relative order of CpG variability of different species only show a little difference with the result using threshold 3σ (Table S2). So choices of the threshold value have small influence on the result.

#### Multi-scale entropy analysis

To quantify the heterogeneity of the CpG distribution, we calculated the multi-scale entropy[2] of the CpG density.

For a data series $X=\{x_1,\ldots,x_i,\ldots,x_N\}$ of length $N$, its m-length vectors are

$$u_m(i)=\{x_i,x_{i+1},\ldots,x_{i+m-1}\},\ 0<i<N-m$$

in which $n^m{}_i(r)$ is defined as the number of m-length vectors $u_m(j)$ that satisfies $d(u_m(i)\ ,\ u_m(j))<r$. Here $d(u_m(i)\ ,\ u_m(j))$ is the distance between two vectors defined as the maximum absolute difference between their components:

$$d(u_m(i),\ u_m(j))=max\{|x_{i+k}-x_{j+k}|:\ 0\leq k\leq m-1\}.$$

Multi-scale entropy (SE) of the data series $X$ is then defined as:

$$SE(m,r,N) = ln\frac{\sum_{i=1}^{N-m} n_i^m}{\sum_{i=1}^{N-m} n_i^{m+1}}$$

*m* and *r* are chosen in this study to be 2 and 1.5 following the reference[2] Costa, et al. (2005).

A higher value of SE indicates a more heterogeneous distribution of *X*. The data series *X* needs to be normalized so that its heterogeneity rather than its integral fluctuation is quantified. To quantify the property of the data series at different length scales, the data can be averaged at different window sizes to yield differently coarse-grained data series, and the multi-scale entropy of each coarse-grained data series can then be calculated.

We applied the Multi-scale entropy analysis to CpG densities of species to describe the CpG distribution in a scale-continuous way. A higher value of multi-scale entropy(SE) corresponds to a more heterogeneous distribution along the genome.

The trends of multi-scale entropy at different average lengths differ substantially among species (Figure S2). SEs of plants and invertebrates decrease in a linear way on the semi-logarithmic graph, suggesting their distribution of CpG becomes more uniform with the increasing average length and no specific mosaic CpG distribution exists in their genome. The SEs of fish, amphibians and reptiles also decrease with the increase of length, but much slower than those of plants and invertebrates. In contrast, SEs of mammals decrease slightly at short genome lengths and then increase at lengths over 100 kb, suggesting a heterogeneous distribution of CpG at a large length scale. Like mammals, SEs of birds also increase with genome length, but to a lesser extent than mammals, leading to largely flat SE curves. SEs of some birds slightly decrease at lengths over 1Mb. This result suggests that CpG distributions of mammals are more heterogeneous than those of birds at the mega-base length scale, consistent with mammals having longer CGI-rich and CGI-poor domains.

At average lengths larger than ~100kb, mammals and birds exhibit strongly heterogeneous CpG density distributions, while those of plants and invertebrates exhibit a low mosaicity, similar to the sequence variations composed of white noise.

**Pearson correlation**

Long-range Pearson correlations of the CpG density of different species were analyzed to investigate the scaling properties of the CpG distribution.

The CpG density correlations of higher species decay with the increase of genomic distance in the form of power-law (Figure S3 A-D), whereas in lower species this power law decay only persists to a short distance if it does exist. Such a power law decay of correlation coefficient reflects the scale-free property of the CpG distribution in the higher but not lower species. Although birds and mammals both exhibit power law decays, they differ substantially in that correlation coefficients of the former decrease more rapidly than those of the latter as the genomic distance increases.

**Scatter plot of average CpG density and CpG variability in different species**

To get a clear relation between average CpG density and CpG variability of species, besides 38 model species, we analyzed another 68 species with genomes available at UCSC genome browser. CpG densities of the genomes are first decomposed by Hilbert Huang Transform, then CpG variabilities are calculated as described in Methods.

**Calculation of compartment vector, contact frequency decay of the chromatin and interaction segregation ratio of compartments**

We calculated chromatin compartment vectors based on the PCA of normalized Hi-C contact matrix, following the method[3] in reference Lieberman-aiden et al. (2009).

We grouped chromatin loci according to genomic distance, then calculated the decay of average Hi-C contact frequency with the genomic distance.

Interaction segregation ratio between A/B compartments is defined as contacts of compartments of the same type (A-A, B-B) divided by contacts of compartments of different types (A-B) of normalized Hi-C contact matrix

**Robustness of structure vectors calculated by different matrix factorization methods**

Besides normal NMF, we also factorized Hi-C matrix by balanced nonnegative matrix factorization[4] (BNMF) and Graph regularized nonnegative matrix factorization[5] (GRNMF) to test the robustness of structure vectors (Figure S6).

### Supplementary Tables

Table S1. The genomes of species used including bacteria, plants, invertebrates, fishes, reptiles, mammals and birds. Genomes of species were retrieved from databases at UCSC and NCBI.

| **Species name** | **genome name** |
|---|---|
| elephant | loxAfr3 |
| gorilla | gorgor5 |
| human | hg19 |
| microbat | myoluc2 |
| mouse | mm10 |
| rat | rn6 |
| Naked_mole_rat | hetgla2 |
| platypus | ornana2 |
| cow | bosTau8 |
| dog | canfam3 |
| manatee | triMan1 |
| pig | susScr11 |
| rabbit | oryCun2 |
| Brown_kiwi | aptMan1 |
| budgerigar | melUnd1 |
| Golden_eagle | aquChr2 |
| Medium_ground_finch | geoFor1 |
| turkey | melGal5 |
| chicken | galgal |
| Zebra_finch | taeGut2 |
| lizard | anocar2 |
| Painted_turtle | chrPic1 |
| X_tropicalis | xenTro9 |
| African_clawed_frog | xenLae2 |
| medaka | oryLat2 |
| Nile_tilapia | orenil2 |
| zebrafish | danrer11 |
| fugu | fr3 |
| tetraodon | tetNig2 |
| lancelet | braFlo1 |
| coelacanth | latCha1 |
| S_purpuratus | strPur2 |
| C_elegans | ce11 |
| D_melanogaster | dm6 |
| S_cerevisiae | sacCer3 |

| | |
|---|---|
| rice | |
| A_thaliana | |
| E_coli | |
| alligator | allMis1 |

Table S2. Comparison of the CpG Variability of species calculated using threshold 2σ and 3σ respectively. The relative order of CpG Variability of different species only shows a little difference.

| Species name | Variability calculated using 2σ | Species name | Variability calculated using 3σ |
|---|---|---|---|
| S_cerevisiae | 3.14048 | S_cerevisiae | 4.094606 |
| E_coli | 3.424367 | E_coli | 4.239483 |
| D_melanogaster | 3.476303 | D_melanogaster | 4.311576 |
| C_elegans | 3.674797 | S_purpuratus | 4.468017 |
| lancelet | 3.728408 | C_elegans | 4.505607 |
| S_purpuratus | 3.743664 | lancelet | 4.619275 |
| fugu | 3.835789 | fugu | 4.713305 |
| A_thaliana | 3.962551 | A_thaliana | 4.807086 |
| rice | 3.9665 | rice | 4.8397 |
| Tetraodon | 3.989295 | Tetraodon | 5.001771 |
| zebrafish | 4.023454 | zebrafish | 5.025087 |
| African_clawed_frog | 4.112653 | African_clawed_frog | 5.105718 |
| medaka | 4.133568 | medaka | 5.129409 |
| Coelacanth | 4.241702 | Coelacanth | 5.20029 |
| Nile_tilapia | 4.286495 | Nile_tilapia | 5.312186 |
| X_tropicalis | 4.791537 | X_tropicalis | 5.992951 |
| lizard | 5.134185 | lizard | 6.372163 |
| platypus | 5.606699 | platypus | 7.034974 |
| Rabbit | 5.87085 | Rabbit | 7.221184 |
| Painted_turtle | 6.090887 | Painted_turtle | 7.602662 |
| microbat | 6.340634 | microbat | 7.869653 |
| Minke_whale | 6.4508 | Minke_whale | 8.0911 |
| rat | 7.110519 | Manatee | 8.755252 |
| pig | 7.30364 | rat | 8.769757 |
| Manatee | 7.379008 | pig | 8.986595 |
| cow | 7.608479 | elephant | 9.07507 |
| elephant | 7.665834 | cow | 9.323523 |
| Gorilla | 7.773693 | human | 9.331596 |
| human | 7.82235 | Gorilla | 9.507512 |
| alligator | 7.95491 | alligator | 9.564039 |
| mouse | 8.166145 | mouse | 10.076576 |
| Brown_kiwi | 8.320886 | Brown_kiwi | 10.20329 |
| Turkey | 8.710931 | Turkey | 10.4812 |
| dog | 8.84555 | Goldern_eagle | 10.65618 |
| Goldern_eagle | 8.959614 | dog | 10.76569 |
| naked_mole_rat | 9.918659 | naked_mole_rat | 11.95753 |
| Medium_ground_finch | 10.16078 | Medium_ground_finch | 12.01873 |
| Budgerigar | 11.06784 | Budgerigar | 13.19898 |

| | | | |
|---|---|---|---|
| Zebra_finch | 11.93517 | Zebra_finch | 14.12514 |
| chicken | 12.13994 | chicken | 14.325685 |

Table S3. Comparison of CpG variability of polar bear and brown bear. CpG variability of polar bear is smaller than the one of brown bear. The assembly level of two genomes is in scaffold. Only several longest scaffolds were analyzed.

| Scaffold ID | CpG Variability | Scaffold Length(kb) |
|---|---|---|
| polarbear_KK498507.1 | 6.235433713 | 4593 |
| polarbear_KK498510.1 | 7.423103699 | 4873 |
| polarbear_KK498511.1 | 6.238884954 | 5261 |
| polarbear_KK498513.1 | 7.065521806 | 40115 |
| polarbear_KK498515.1 | 7.871023126 | 12716 |
| brownbear_KZ986268.1 | 9.342643186 | 92499 |
| brownbear _KZ986269.1 | 9.359353295 | 90872 |
| brownbear _KZ986270.1 | 8.957848845 | 84388 |
| brownbear _KZ986271.1 | 9.533290518 | 79539 |
| brownbear _KZ986272.1 | 9.580195044 | 76399 |
| brownbear _KZ986273.1 | 8.424618173 | 67437 |
| brownbear _KZ986274.1 | 9.090982869 | 58486 |
| brownbear _KZ986275.1 | 9.108413252 | 57860 |
| brownbear _KZ986276.1 | 9.662980646 | 56669 |
| brownbear _KZ986277.1 | 9.627856936 | 53000 |
| brownbear _KZ986278.1 | 9.348755007 | 52083 |
| brownbear _KZ986279.1 | 9.172397596 | 51855 |

Table S4. CpG variability of fish from regions of low temperature such as the Antarctic region, most of which are smaller than the ones of normal fish (CpG Variability ~ 5).

*Assembled in scaffold level, scaffolds longer than 30kb were jointed for analyzing.

| Fish Name | CpG Variability |
|---|---|
| Benthosema_glaciale* | 4.1 |
| Arctogadus_glacialis* | 4.1 |
| Gadus_morhua | 4.5 |
| Labrus_bergylta | 4.6 |
| Chaenocephalus_aceratus* | 4.7 |
| Notothenia_coriiceps | 4.8 |
| Esox_lucius | 5.2 |

Table S5. Source of Hi-C datasets.

| Species name | Source | accession number |
|---|---|---|
| human | Schmitt et al.2016 | GSE87112 |
| mouse | Schwarzer et al.2017 | GSE93431 |
| chicken | Brossas et al.,2020 | GSE153566 |
| zebrafish | Yang et al.,2020 | GSE134055 |
| yeast | Kim et al.,2017 | GSE88952 |
| athaliana | Grob et al.2014 | GSE55960 |
| S.acidocaldarius | Takemata et al.2019 | GSE128063 |
| drosophila | Sharnouby et al.2017 | GSE85503 |
| rice | Liu et al.2017 | SRP093806 |

Table S6. URL of RNA-seq datasets

| Species name | URL |
|---|---|
| human | https://zenodo.org/record/838734 |
| rice | https://expression.ic4r.org |

Table S7. Cell types used in performing Wavelet Transform of structure vector and CpG density of species.

| Species name | Cell type |
|---|---|
| human | liver, IMR90 |
| mouse | liver |
| chicken | DT40 |
| zebrafish | muscle |
| athaliana | seedling |
| S.acidocaldarius | / |
| yeast | / |

Table S8. Max absolute value of Pearson correlation of compartment vector and structure vector of species.

| Species | Cell type | Pearson correlation |
|---|---|---|
| chicken | DT40 | 0.8773 |
| mouse | liver | 0.9067 |
| human | IMR90 | 0.8809 |
| zebrafish | muscle | 0.9646 |
| A.thaliana | seedling | 0.8237 |
| yeast | / | 0.9011 |
| S.acidocaldarius | / | 0.976 |

Table S9. Pearson correlation of CpG-rich/poor domain index and A/B compartment index.

| Species | Cell type | Pearson correlation |
|---|---|---|
| human | IMR90 | 0.4969 |
| mouse | liver | 0.3459 |
| chicken | DT40 | 0.5379 |
| zebrafish | muscle | 0.0617 |
| A.thaliana | seedling | 0.2023 |
| yeast | / | -0.0192 |

Table S10. Interaction segregation ratio of A/B compartments of species.

| Species | Segregation ratio |
|---|---|
| S.acidocaldarius | 1.1435 |
| yeast | 1.6231 |
| A.thaliana | 1.4186 |
| zebrafish | 1.3572 |
| drosophila | 1.315 |
| chicken | 1.4974 |
| human | 1.7392 |
| mouse | 2.0888 |

#### Supplementary Figures

A

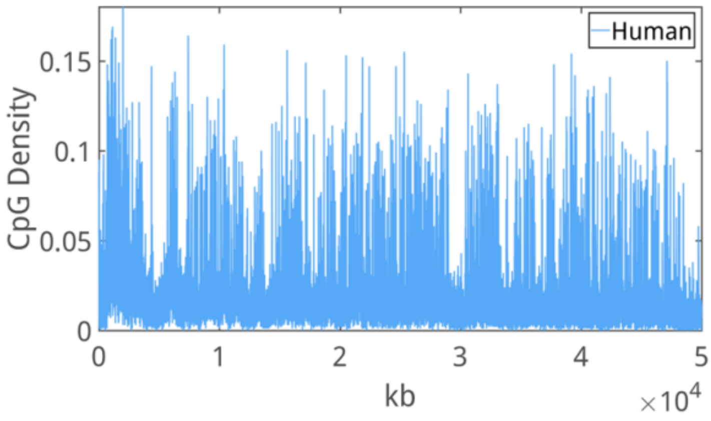


B

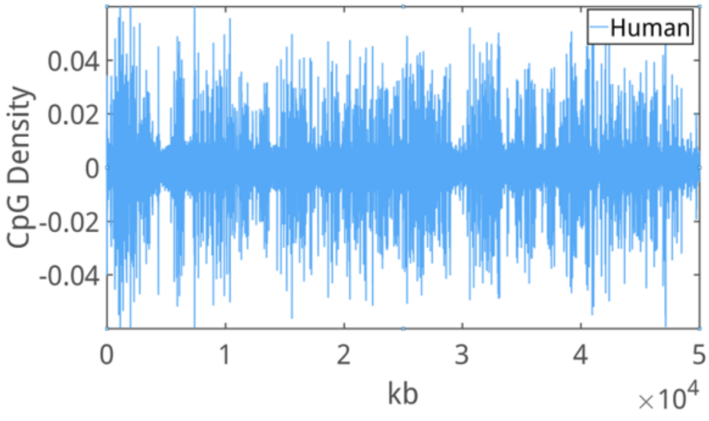


C

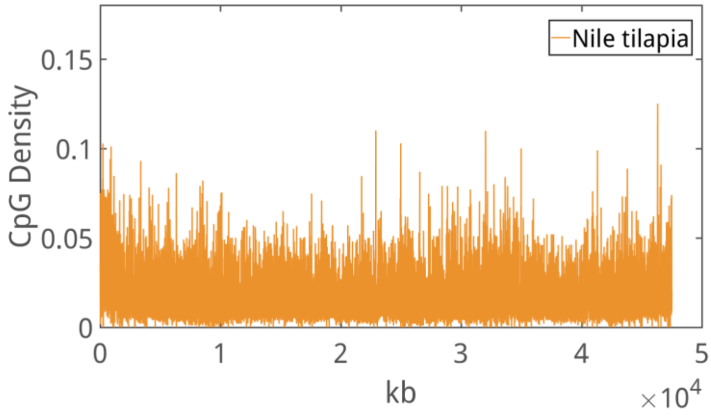


D

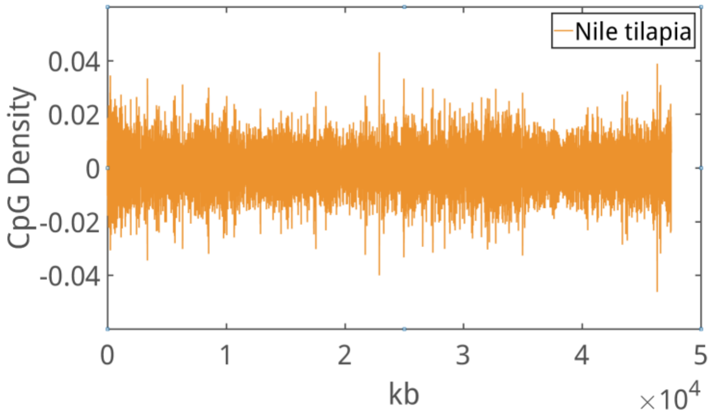


Figure S1. Representative curves of (A, C) CpG density and (B, D) decomposed series of CpG density along the genomes of human and Nile tilapia. There are sharp peaks in the decomposed series of CpG density of human, while not in the ones of Nile tilapia. Accordingly, the CpG Variability of human is larger than the one of Nile tilapia.

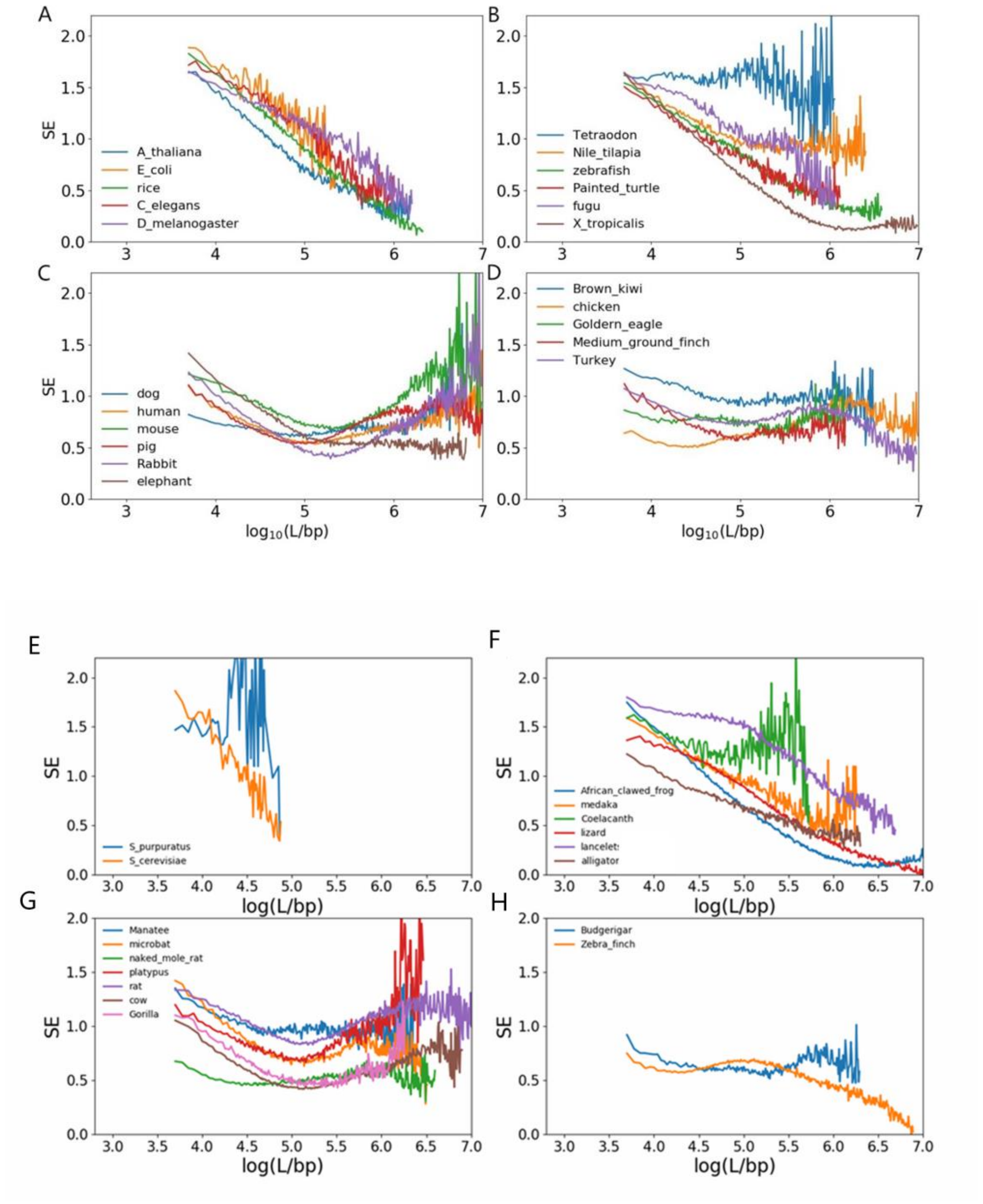


Figure S2. Multi-scale Entropy (SE) of CpG density of species. (A,E) bacteria, invertebrates and plants, (B,F) fishes, amphibians and reptiles, (C,G) mammals, (D,H) birds.

A

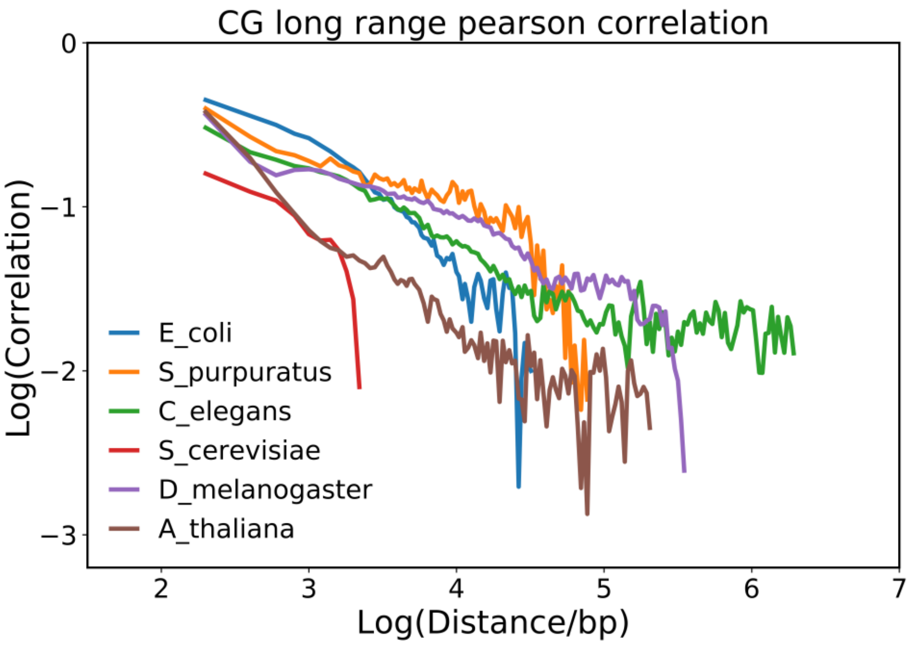

CG long range pearson correlation
Log(Correlation)
Log(Distance/bp)
0
−1
−2
−3
2
3
4
5
6
7
E_coli
S_purpuratus
C_elegans
S_cerevisiae
D_melanogaster
A_thaliana


B

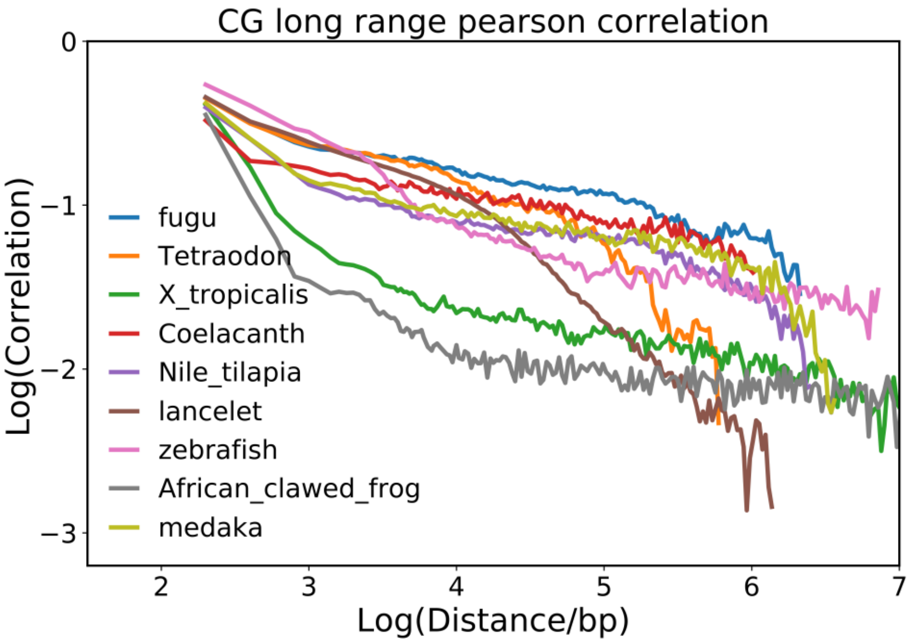

CG long range pearson correlation
Log(Correlation)
Log(Distance/bp)
0
−1
−2
−3
2
3
4
5
6
7
fugu
Tetraodon
X_tropicalis
Coelacanth
Nile_tilapia
lancelet
zebrafish
African_clawed_frog
medaka

C

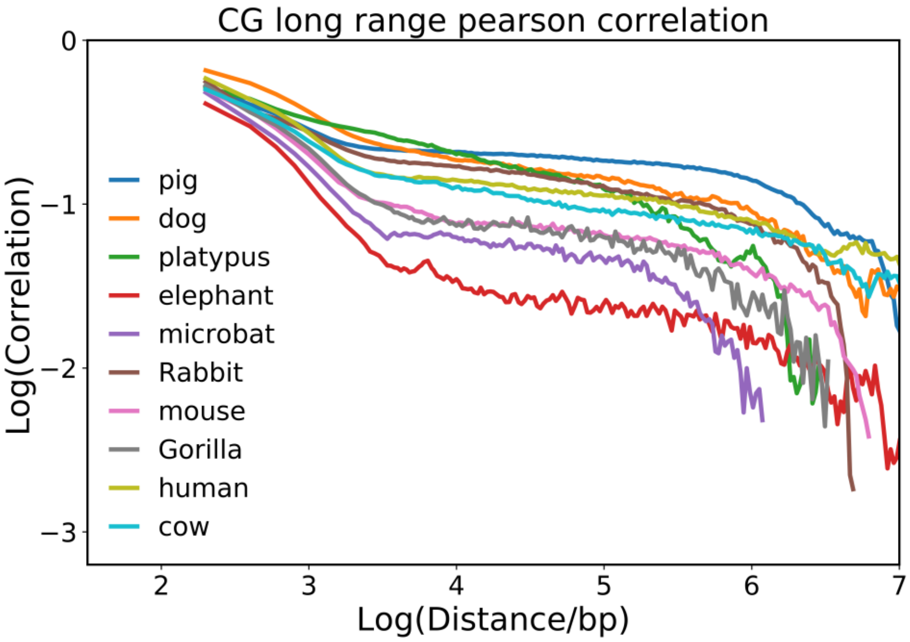
CG long range pearson correlation
0
−1
−2
−3
Log(Correlation)
2
3
4
5
6
7
Log(Distance/bp)
pig
dog
platypus
elephant
microbat
Rabbit
mouse
Gorilla
human
cow

D

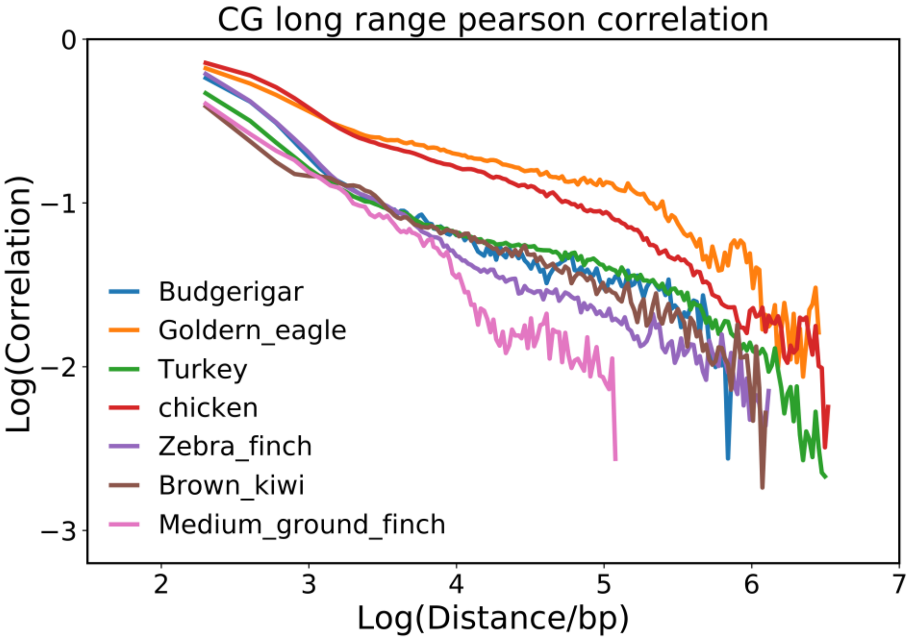
CG long range pearson correlation
0
−1
−2
−3
Log(Correlation)
2
3
4
5
6
7
Log(Distance/bp)
Budgerigar
Goldern_eagle
Turkey
chicken
Zebra_finch
Brown_kiwi
Medium_ground_finch

E

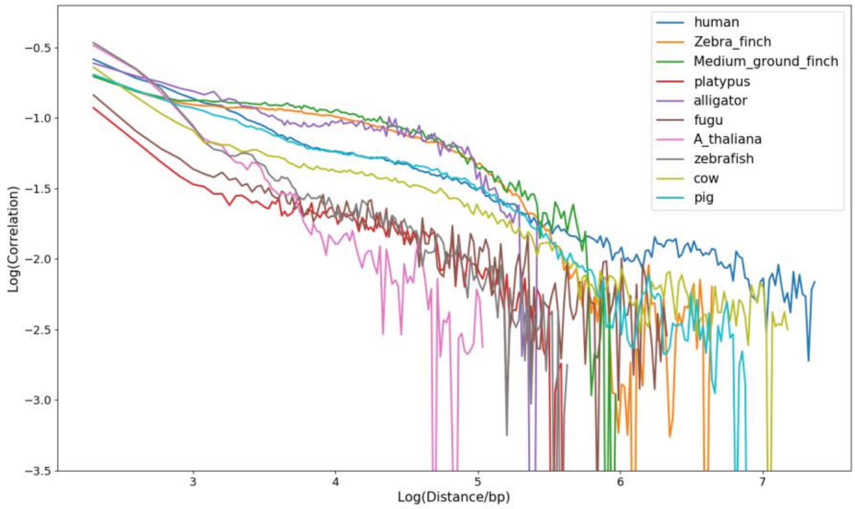


F

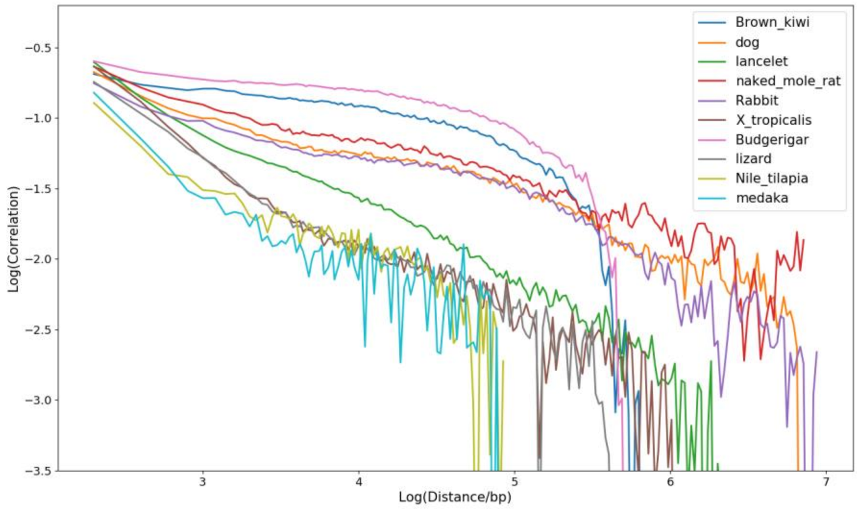


Figure S3.
(A-D) Long-range Pearson correlation of CpG density distribution of (A) bacteria, invertebrates and plants, (B)fishes and amphibians,(C) mammals, (D)birds.
(E)(F)Long-range Pearson correlation of CpA density distribution of species.

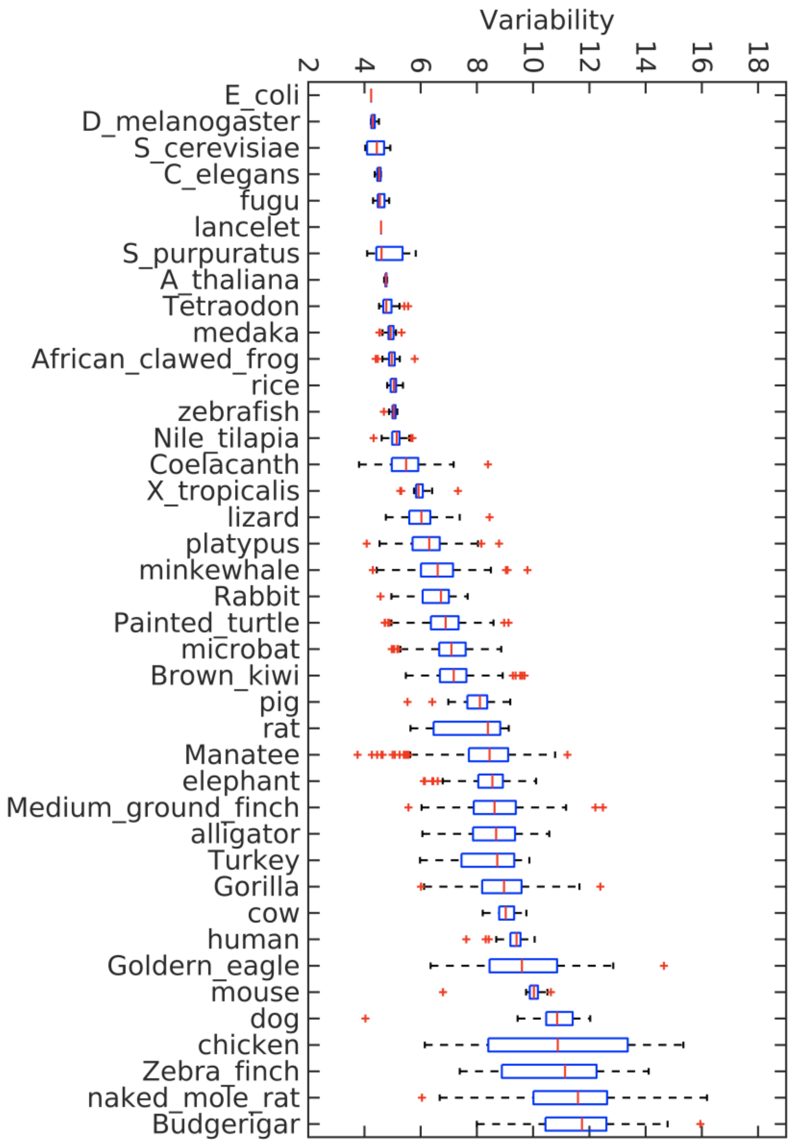


Figure S4. Boxplot of variabilities of the CpG density of chromosomes of different species.

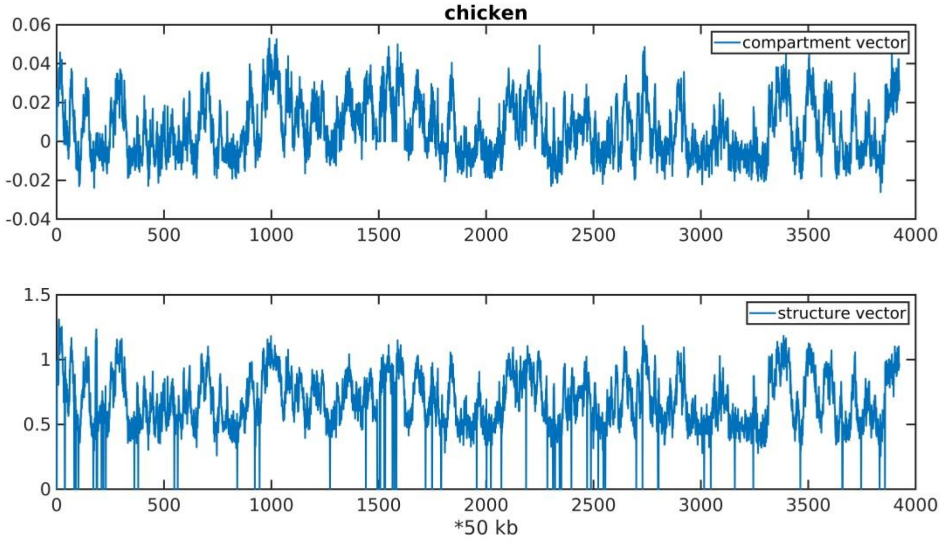


Figure S5. Compartment vector and structure vector along chr1 of chicken.

A

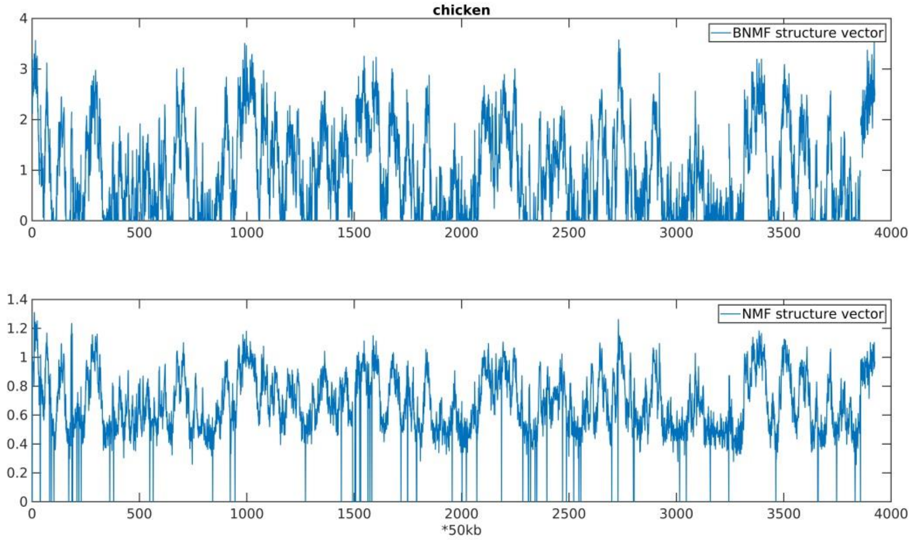
chicken
BNMF structure vector
NMF structure vector
*50kb

B

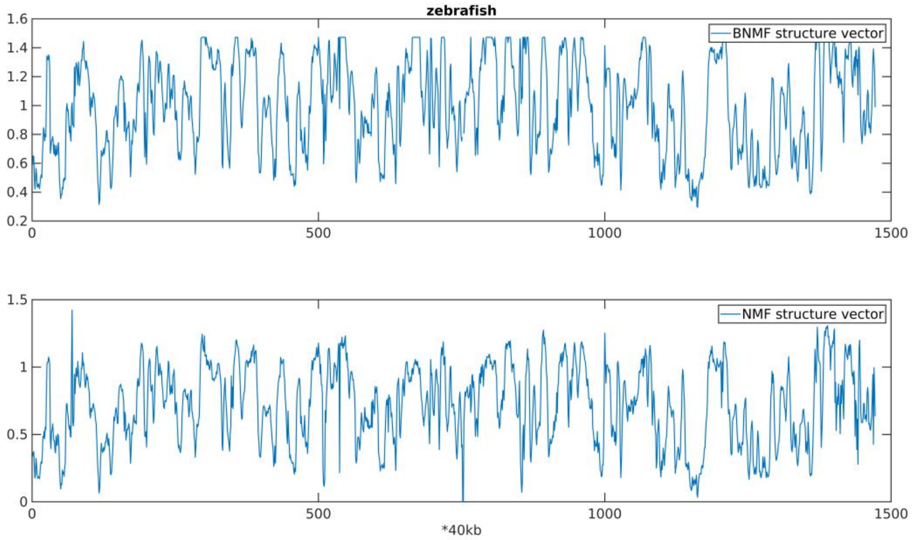
zebrafish
BNMF structure vector
NMF structure vector
*40kb

C

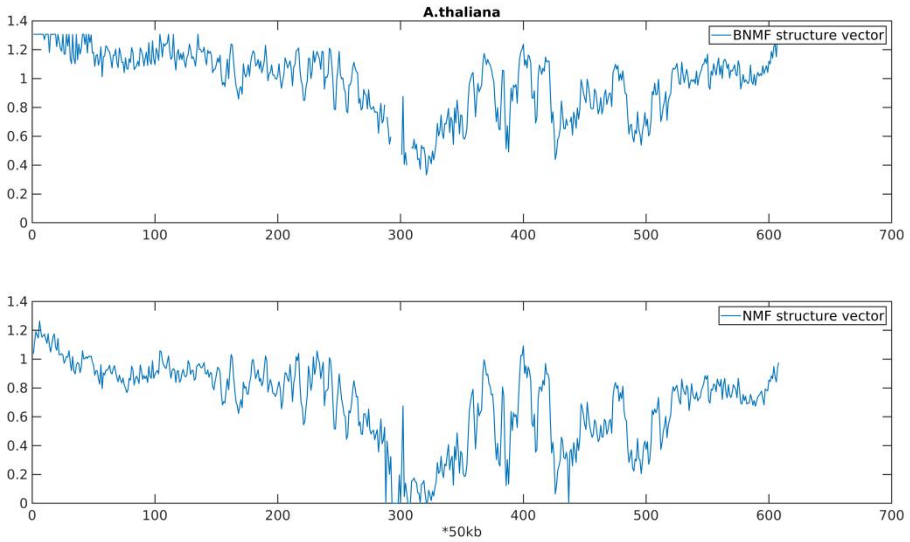

A.thaliana
BNMF structure vector
NMF structure vector
*50kb


**D**

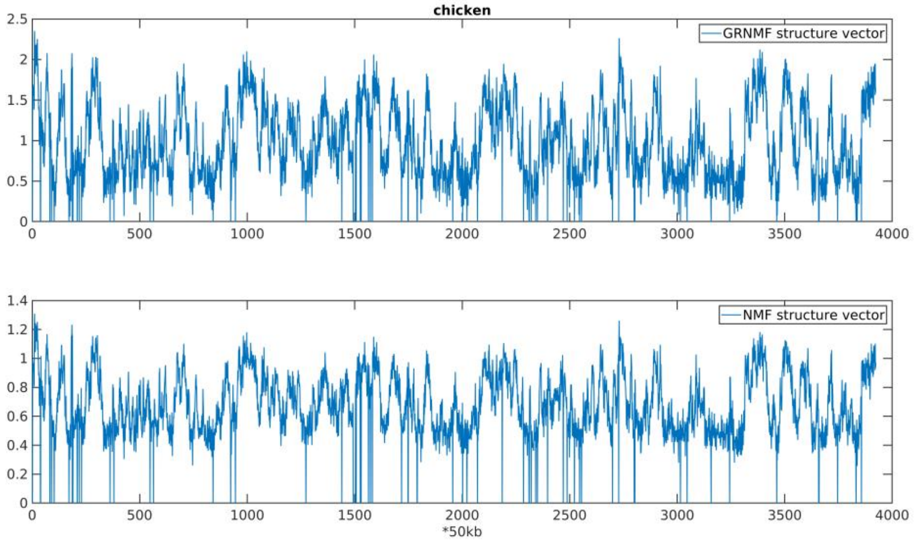

chicken
GRNMF structure vector
NMF structure vector
*50kb

E

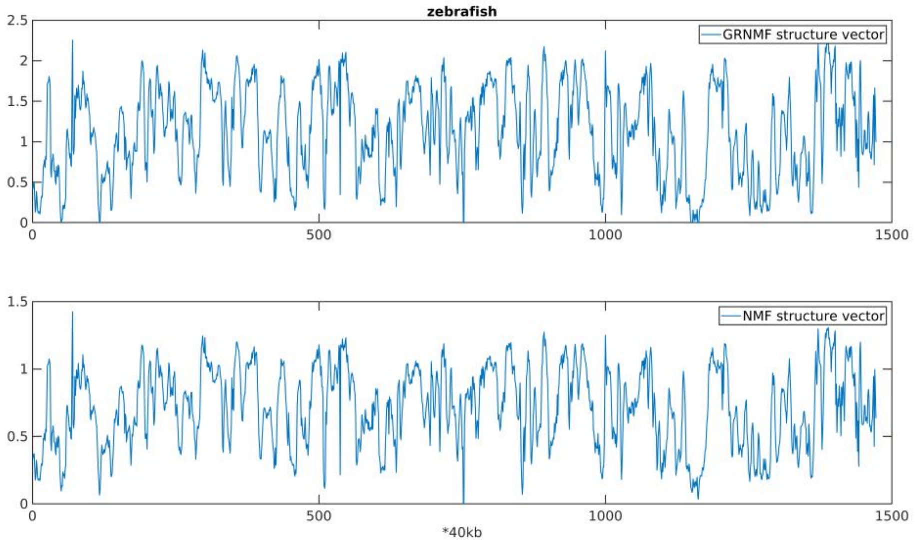


F

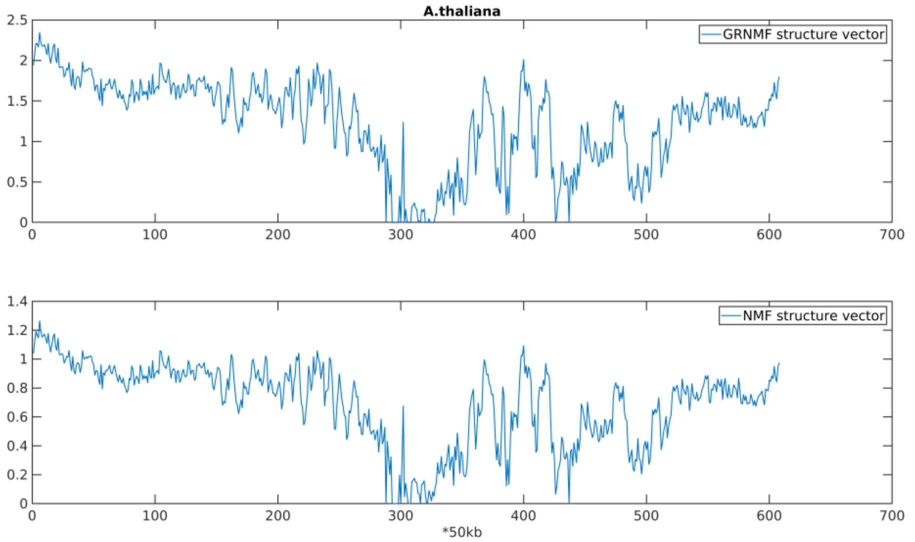


Figure S6. (A-C) Comparison of structure vectors factorized by BNMF and NMF in chicken, zebrafish, A.thaliana. (D-F) Comparison of structure vectors factorized by GRNMF and NMF in chicken, zebrafish, A.thaliana.

A

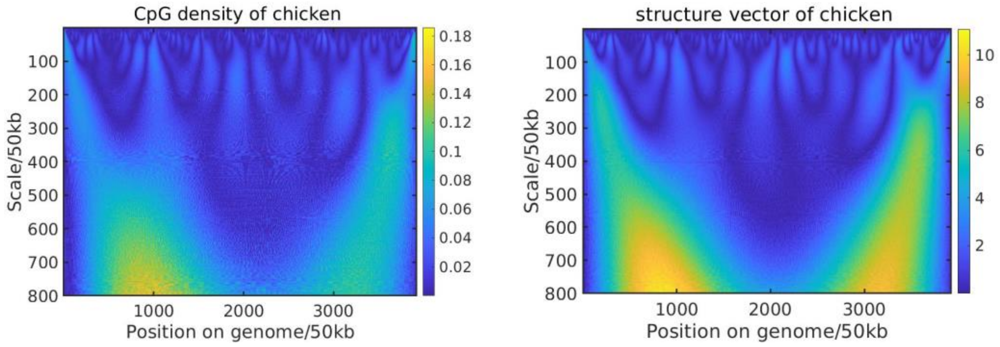


B

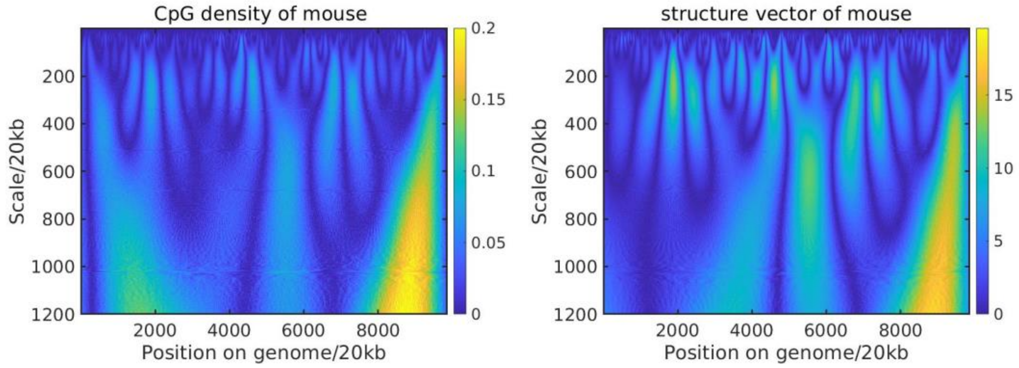


C

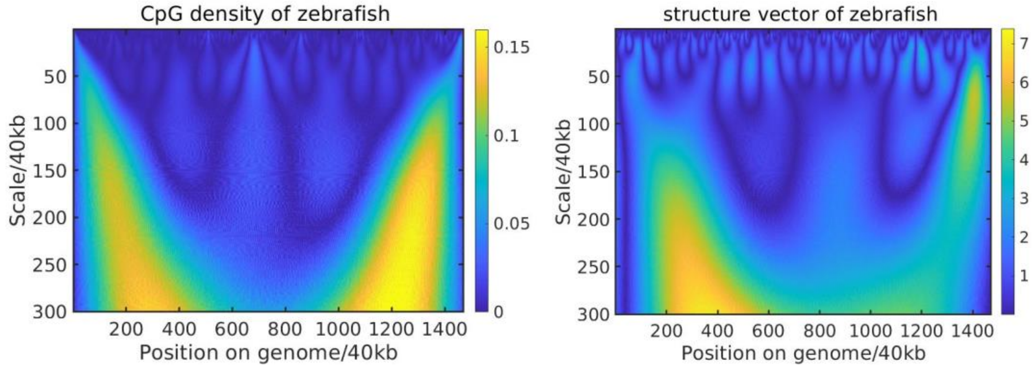


D

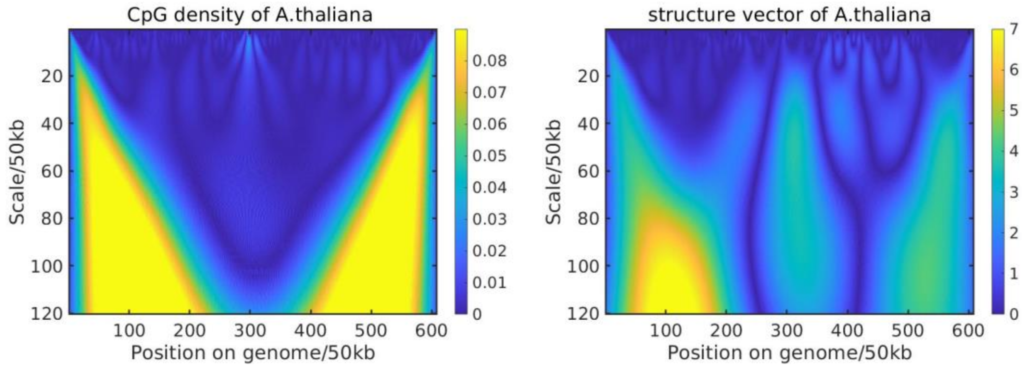


Figure S7. Wavelet Transform of CpG density and structure vector (W1) along the genome of (A) chicken, (B) mouse, (C) zebrafish, (D) A.thaliana.

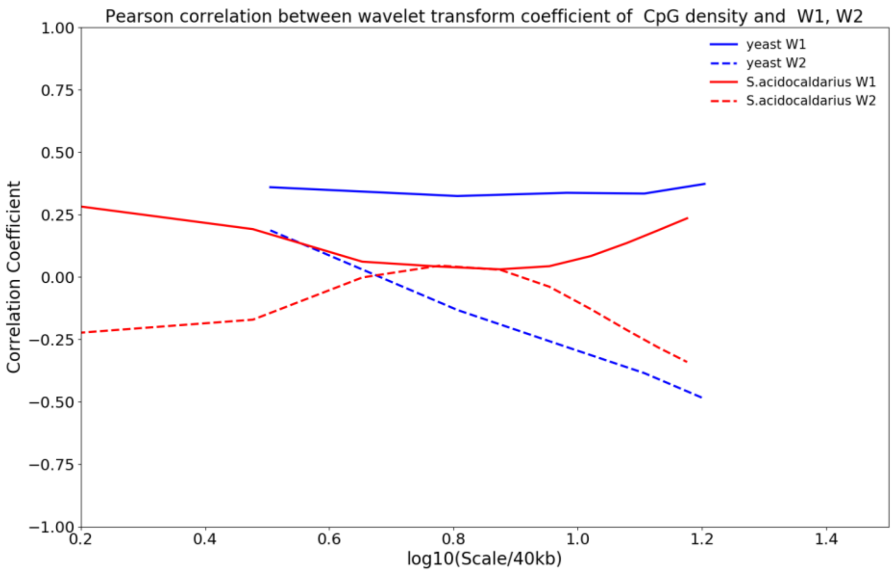


Figure S8. Pearson correlation of Wavelet Transform coefficient of structure vectors (i.e. W1 and W2) and CpG density of yeast and S.acidocaldarius.

A

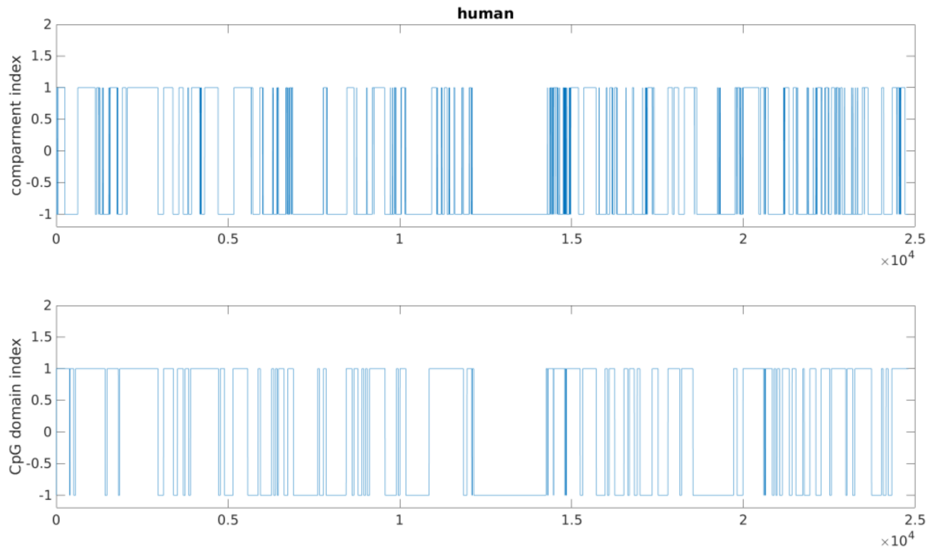


B

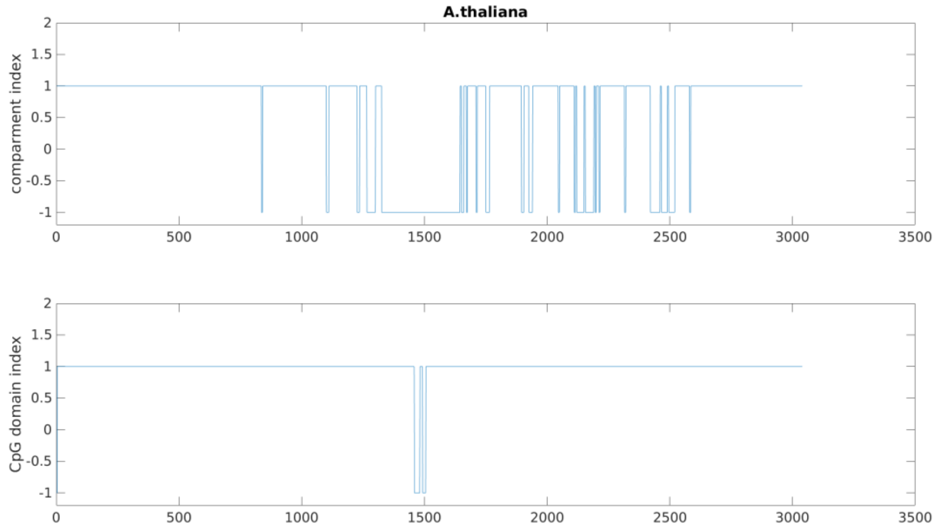


Figure S9. Comparison of CpG-rich/poor domain and A/B compartments of (A)human and (B)A.thaliana. CpG domain index is 1 for CpG-rich domain and -1 for CpG-poor domain. Compartment index is 1 for compartment A and -1 for compartment B.

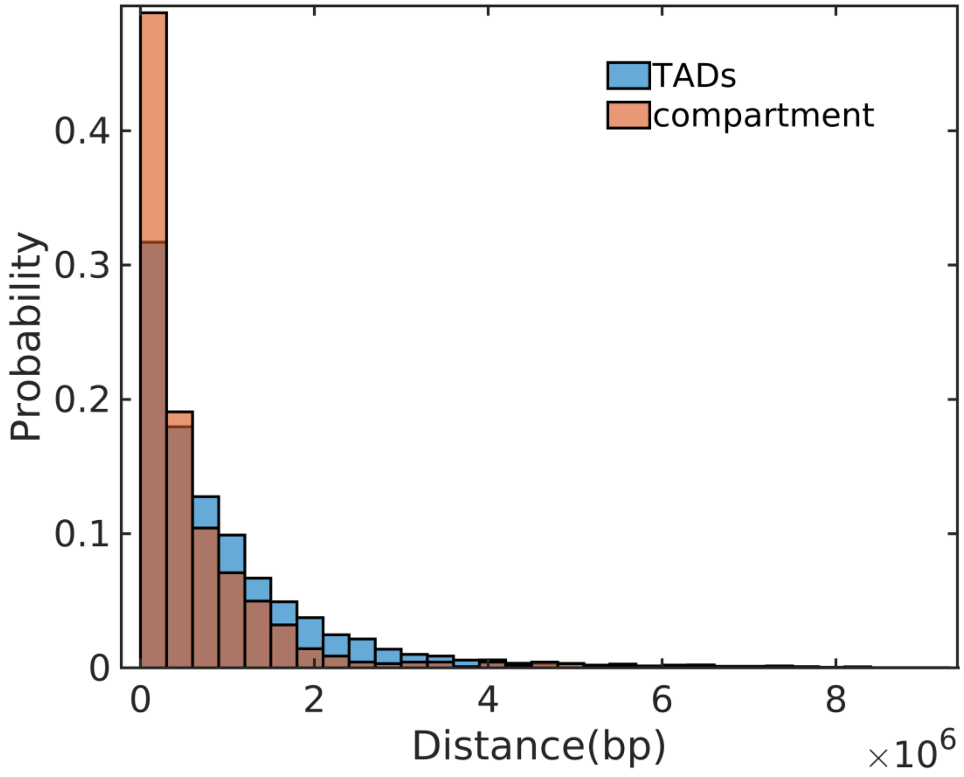


Figure S10. Distance distribution of TAD boundaries to CpG-rich/poor domain boundaries and compartments boundaries to CpG-rich/poor domain boundaries in human.

A

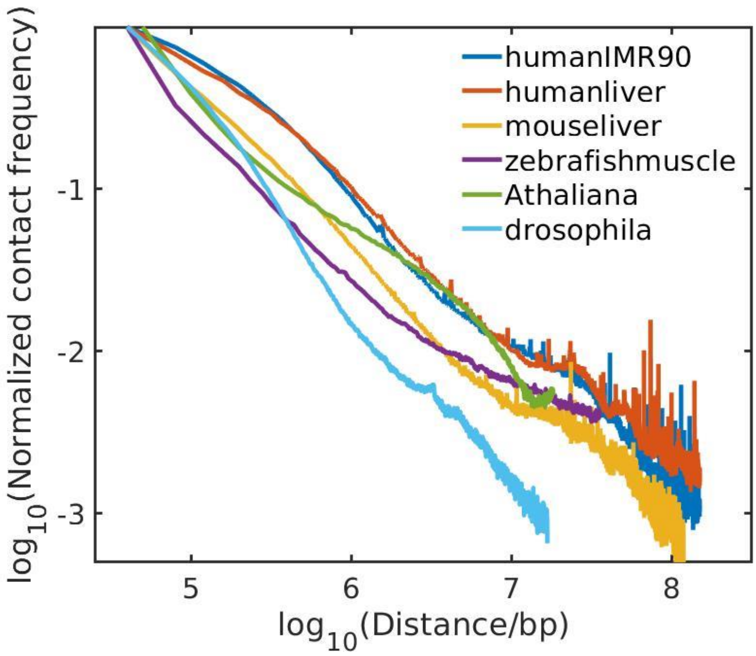


B

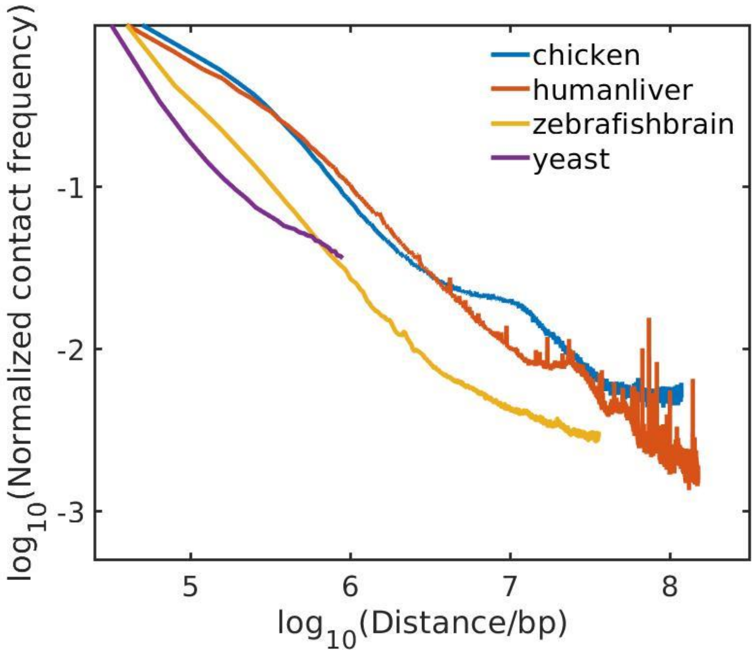


Figure S11. The dependence of contact frequency of chromatin on the genome distance for various species.

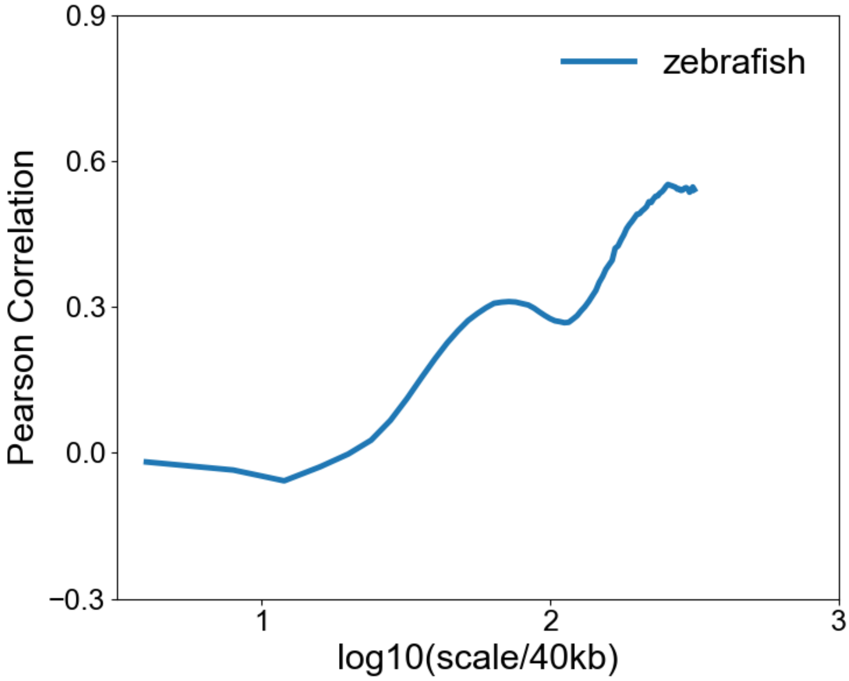


Figure S12. Pearson correlation of Wavelet transform coefficient of CpG density and expression level at multi-length scales of zebrafish.

A

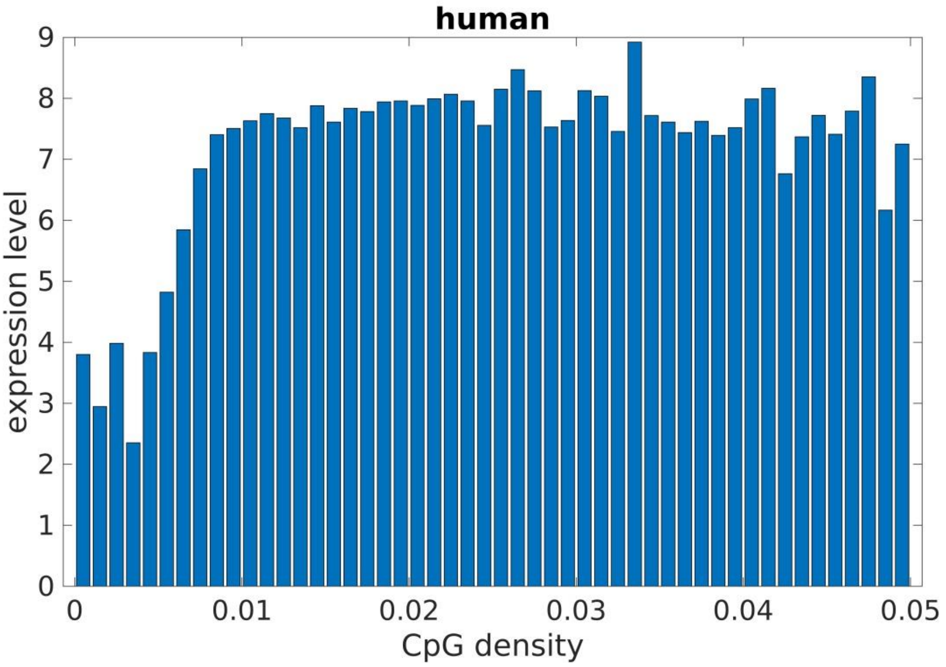

B

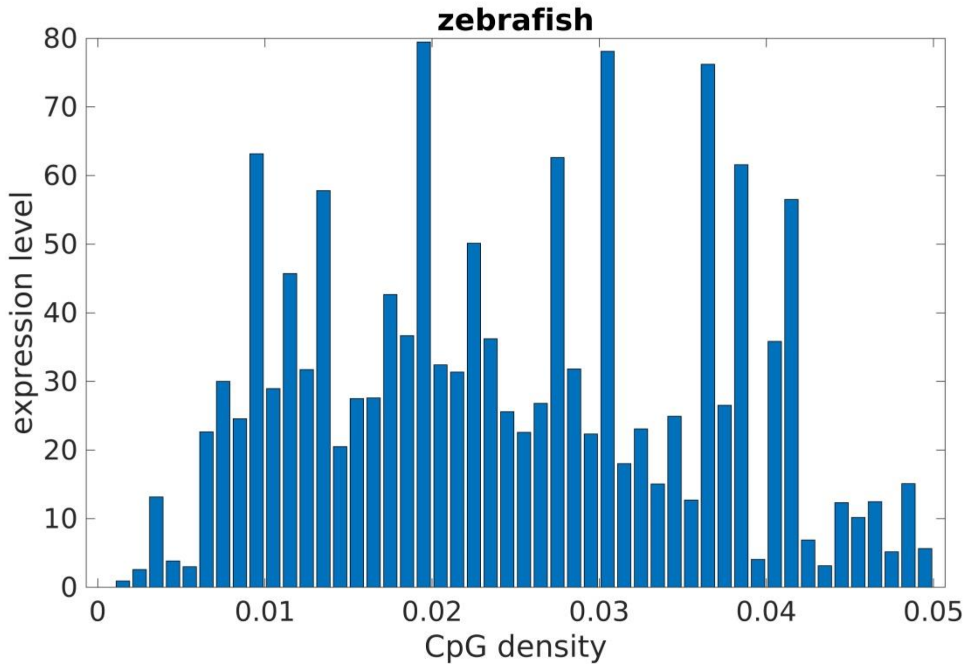


C

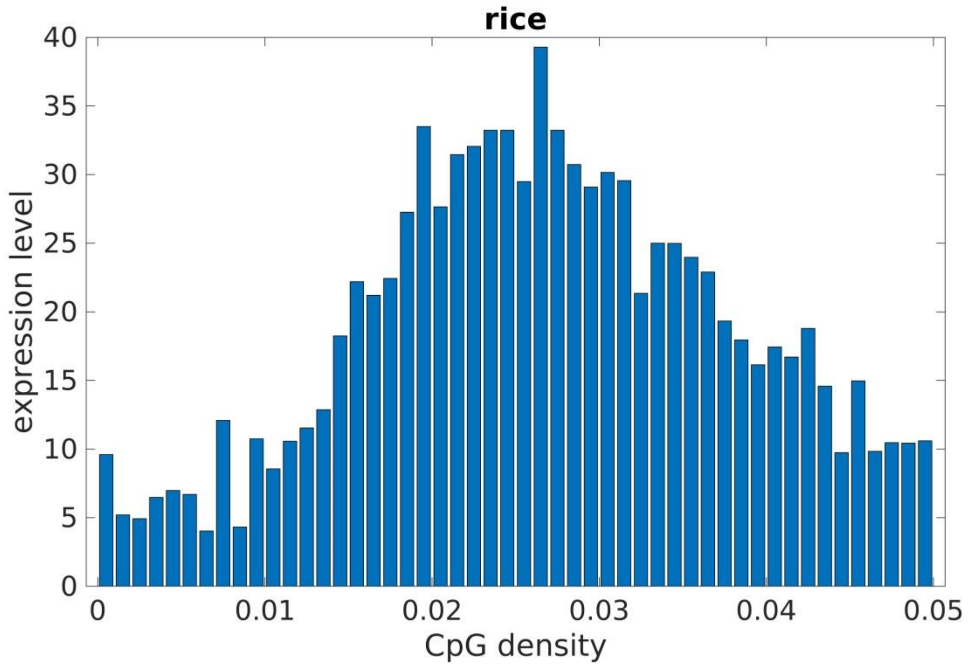


Figure S13. Expression level of genes from groups with different CpG densities in human(A), zebrafish(B), rice(C). There is a general increasing trend of expression level with CpG density of genes in human. The positive correlation of expression level and CpG density of genes is higher in human than in zebrafish and in rice.

A

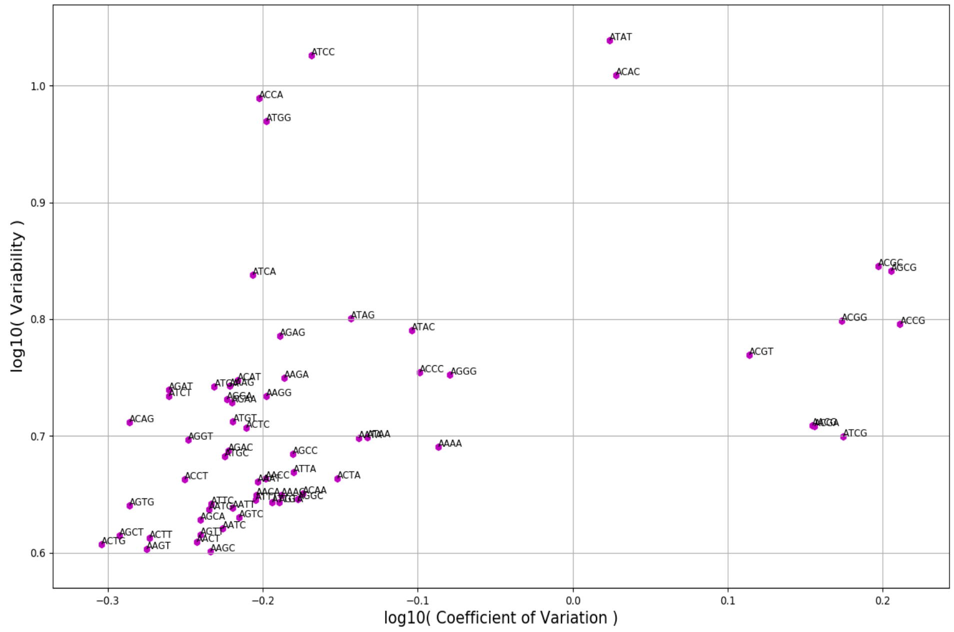
log10( Variability )
log10( Coefficient of Variation )
1.0
0.9
0.8
0.7
0.6
−0.3
−0.2
−0.1
0.0
0.1
0.2
ATAT
ATCC
ACAC
ACCA
ATGG
ACGC
AGCG
ATCA
ATAG
ACGG
ACCG
ATAC
AGAG
ACGT
ACCC
AGGG
AAGA
ACAT
AGAT
ATCT
AAGG
ACAG
ATGT
ACTC
ATCG
AAAA
AGGT
AGAC
AGCC
ATTA
ACCT
ACTA
AGTG
ATTC
AGCA
AGTC
AATC
AGCT
ACTT
ACTG
AAGT
AAGC

B

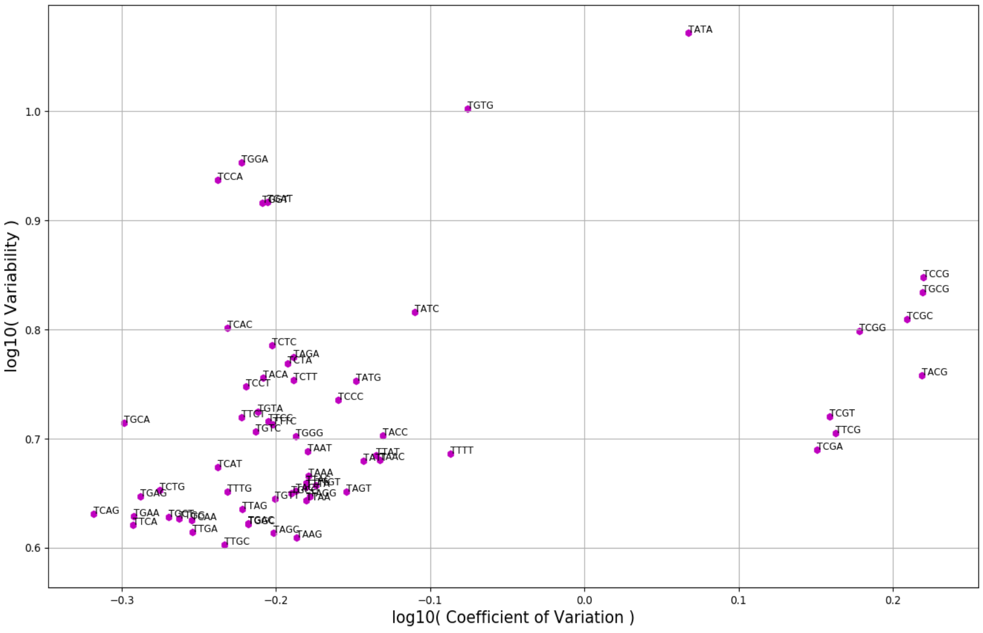
log10( Variability )
log10( Coefficient of Variation )
1.0
0.9
0.8
0.7
0.6
−0.3
−0.2
−0.1
0.0
0.1
0.2
TATA
TGTG
TGGA
TCCA
TCCG
TGCG
TCGC
TCGG
TATC
TCAC
TCTC
TAGA
TCTA
TACA
TCTT
TATG
TCCT
TACG
TCCC
TGTA
TGCA
TCGT
TTCG
TGTC
TGGG
TACC
TCGA
TAAT
TTTT
TCAT
TAAA
TAGT
TCTG
TTTG
TGAG
TTAG
TCAG
TGAA
TTCA
TTGA
TTGC
TAGG
TAAG

C

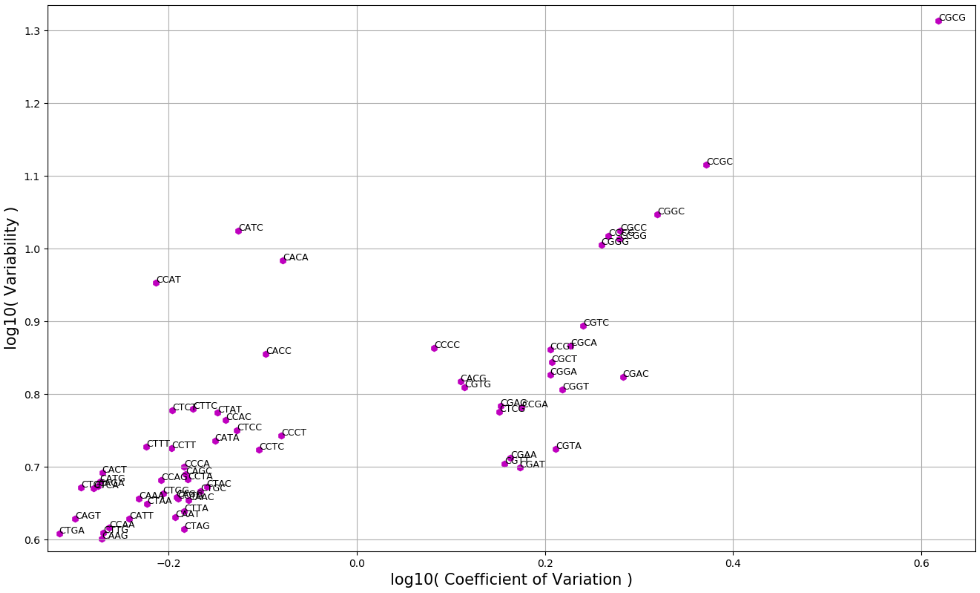


D

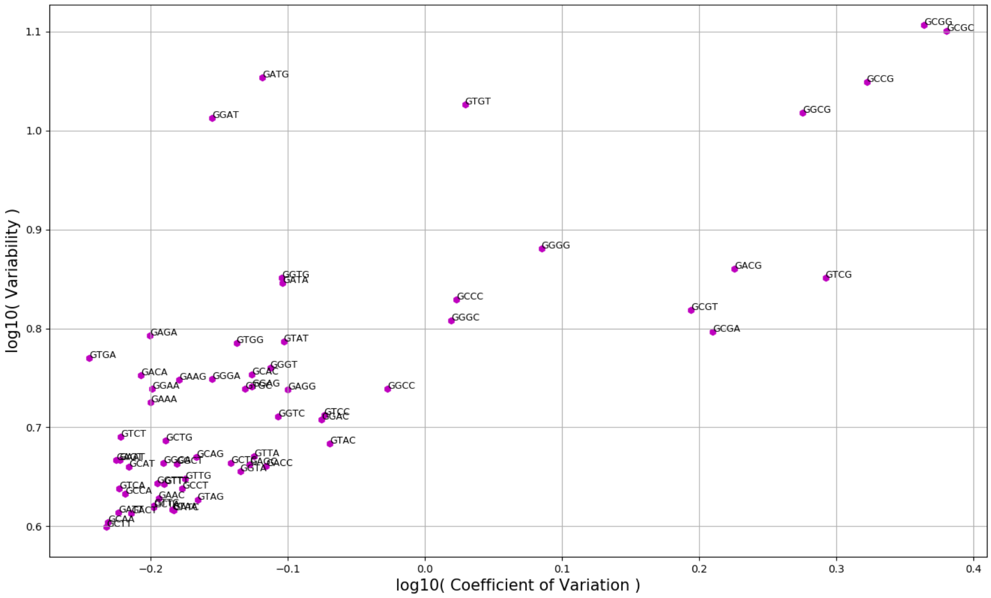


Figure S14. The scatter plot for Variability and CV of 256 tetranucleotides density in human chr1. (A,B,C,D) The scatter plot of tetranucleotides which started with A, T, C and G, respectively.

A

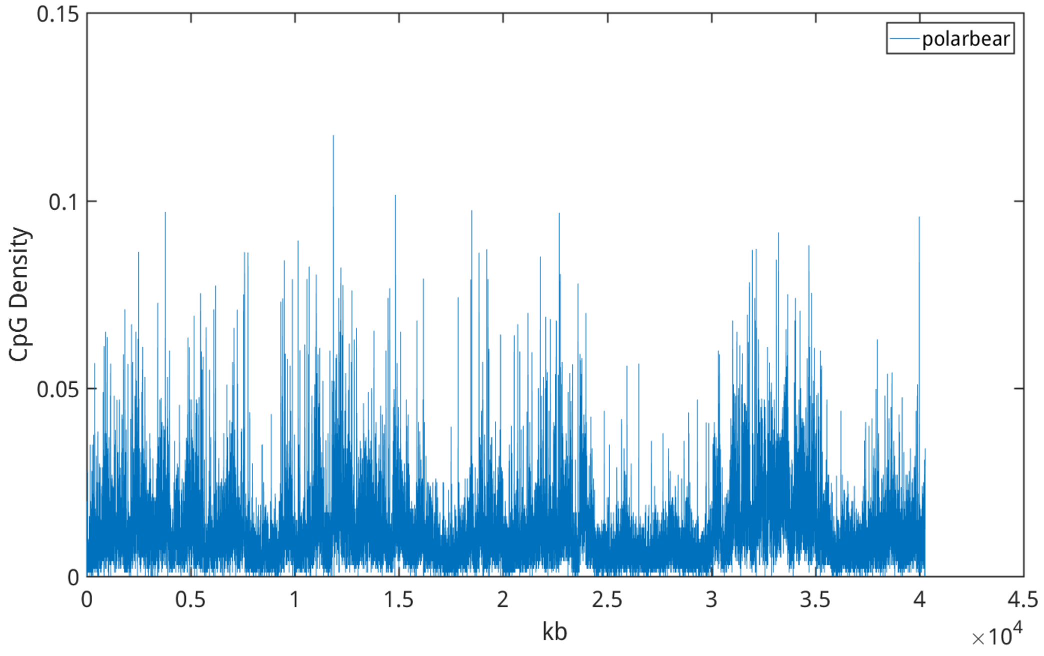


B

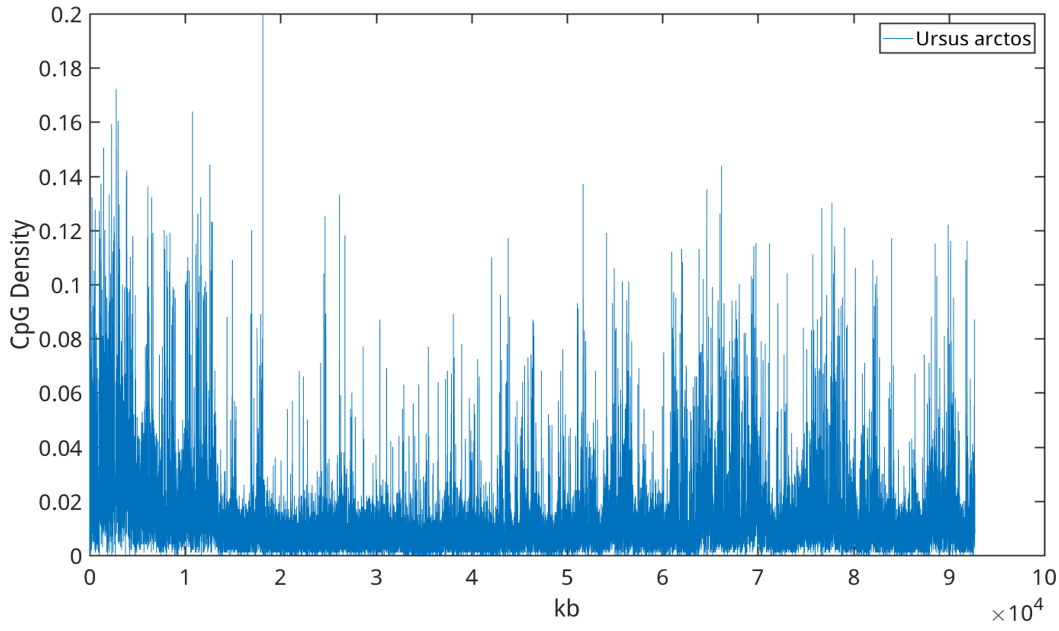


Figure S15. Representative curves of CpG density of (A) polar bear and (B) brown bear along the sequence.

A

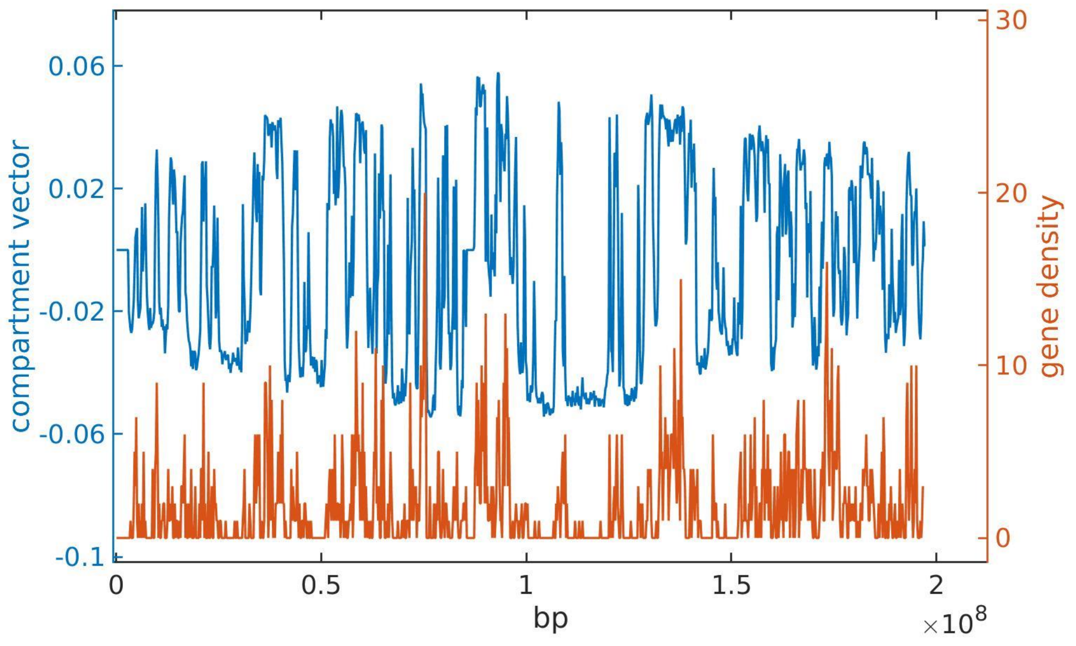

0.06
0.02
-0.02
-0.06
-0.1
compartment vector
30
20
10
0
gene density
0
0.5
1
1.5
2
bp
×10^8


B

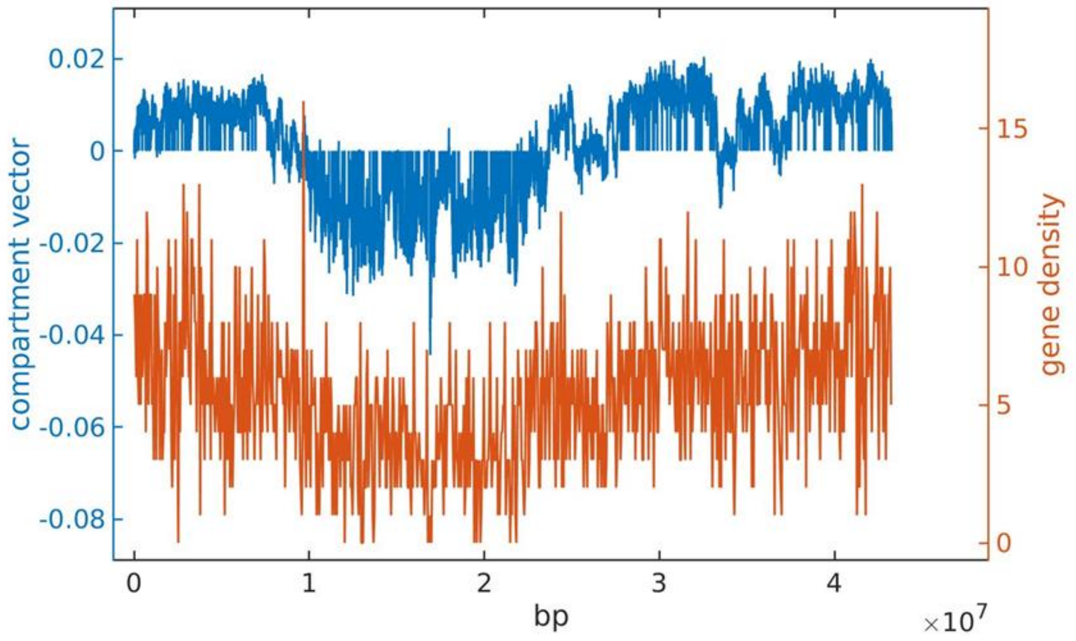

0.02
0
-0.02
-0.04
-0.06
-0.08
compartment vector
15
10
5
0
gene density
0
1
2
3
4
bp
×10^7

C

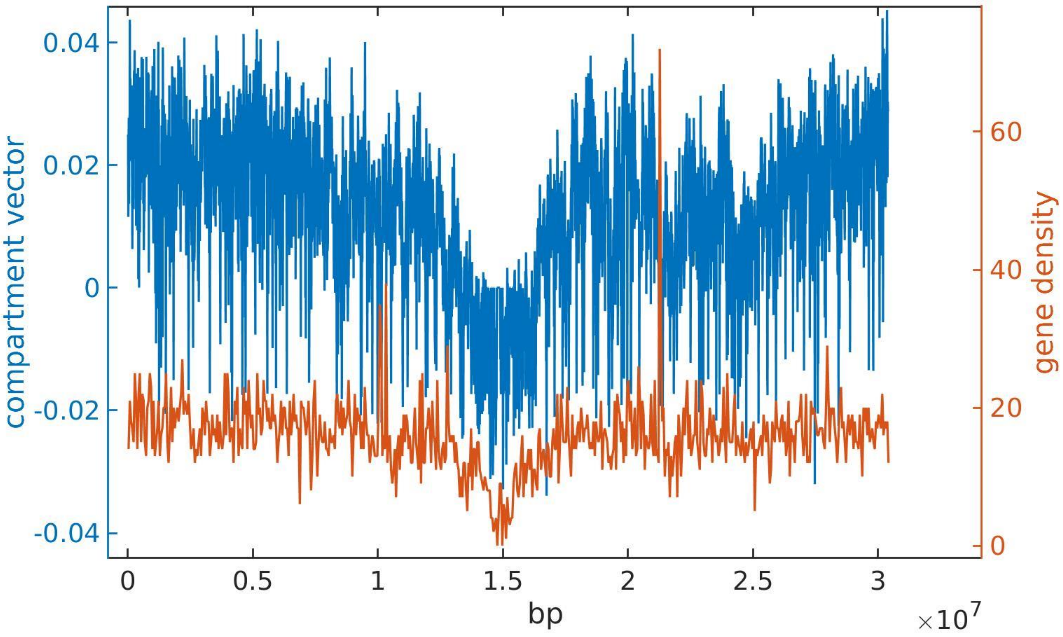


D

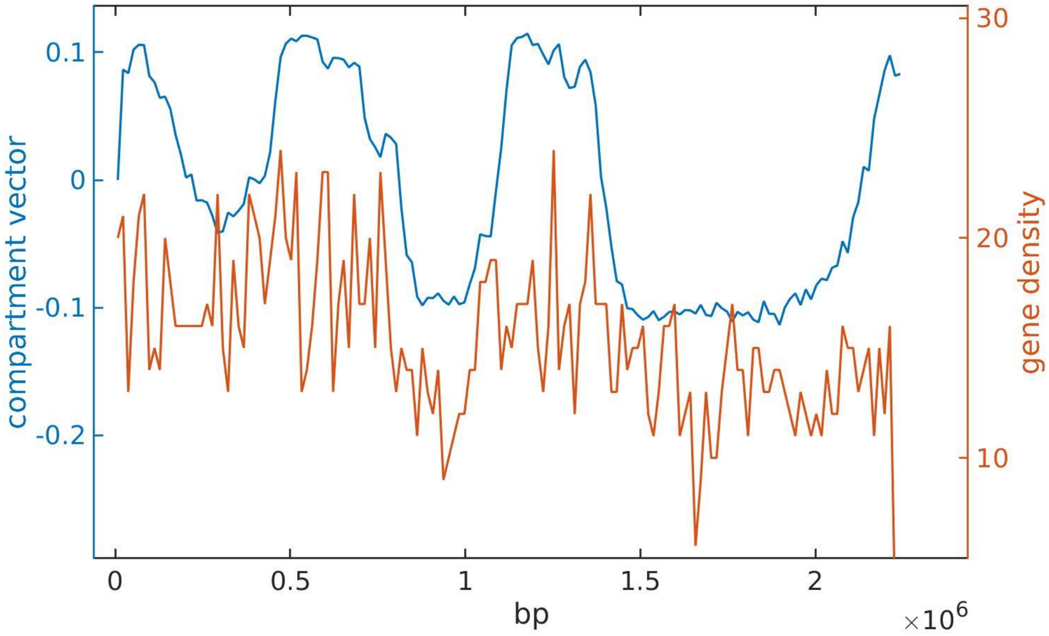


Figure S16. Comparison between chromosomal compartment vector (blue) and gene density (red) distribution along the genomes of (A) mouse, (B) rice, (C) A.thaliana and (D) S.acidocaldarius.
